# Lineage motifs: developmental modules for control of cell type proportions

**DOI:** 10.1101/2023.06.06.543925

**Authors:** Martin Tran, Amjad Askary, Michael B. Elowitz

**Affiliations:** Division of Biology and Biological Engineering, California Institute of Technology, Pasadena, CA 91125, USA; Department of Molecular, Cell and Developmental Biology, University of California Los Angeles, Los Angeles, CA 90095, USA; Howard Hughes Medical Institute, Chevy Chase, MD 20815, USA

**Keywords:** Developmental programs, lineage tree analysis, cell fate correlations, motif analysis, cell type proportions, Pareto optimality, retina development, blastocyst development

## Abstract

In multicellular organisms, cell types must be produced and maintained in appropriate proportions. One way this is achieved is through committed progenitor cells that produce specific sets of descendant cell types. However, cell fate commitment is probabilistic in most contexts, making it difficult to infer progenitor states and understand how they establish overall cell type proportions. Here, we introduce Lineage Motif Analysis (LMA), a method that recursively identifies statistically overrepresented patterns of cell fates on lineage trees as potential signatures of committed progenitor states. Applying LMA to published datasets reveals spatial and temporal organization of cell fate commitment in zebrafish and rat retina and early mouse embryo development. Comparative analysis of vertebrate species suggests that lineage motifs facilitate adaptive evolutionary variation of retinal cell type proportions. LMA thus provides insight into complex developmental processes by decomposing them into simpler underlying modules.

## Introduction

Most tissues comprise multiple specialized cell types that appear in appropriate proportions to support proper tissue-level functions. In many cases, cell type proportions vary spatially within the tissue. For example, the center of the primate retina is cone-dense, allowing for high visual acuity in the fovea, while the periphery is rod-dense, enabling greater sensitivity in low light conditions ^1^. Cell type proportions also vary between species. For instance, the ratio of rod and cone photoreceptors varies depending on the visual needs associated with the lifestyle of each species ^2^. Tissue development thus faces the fundamental challenges of (1) generating cell types in correct proportions, and (2) facilitating spatial and evolutionary changes in those proportions ^3, 4^.

One prevalent mechanism for specifying cell type proportions occurs through developmental programs, which determine the probabilities with which progenitor cells progressively become restricted in their fate potential or competence, and eventually commit to terminal cell fates. In some cases, like the nematode *C. elegans*, the developmental program can be deterministic, producing a stereotyped lineage tree in all individuals ^5^. However, in most other organisms, one cannot infer a general program from any single lineage tree due to variability. For example, in the mammalian retina, individual progenitor cells can give rise to a wide distribution of cell numbers and types with no apparent fixed ratios between different types, prompting investigators to initially suggest a stochastic view of cell fate determination ^6, 7^. However, other studies of terminally dividing progenitors with particular expression patterns provided evidence for consistent cell-intrinsic biases in cell fate decisions ^8–12^. These biases can extend upstream to non-terminal divisions ^10, 13^. Developmental programs can also integrate extrinsic signals, spatial context, developmental time, cell history, and stochastic “noise” with internal progenitor states ^14, 15^. Thus, even in well-studied systems such as the retina, it remains a major challenge to elucidate developmental programs.

Different developmental programs can generate distinct distributions of cell fates on lineage trees. One of the simplest possible developmental programs comprises a multipotent progenitor that can directly and probabilistically differentiate into multiple terminal fates (**Figure 1A**). A system employing such a direct, memoryless program would not exhibit fate correlations between lineally related cells (sisters, cousins, etc.). Alternatively, a more complex program could involve the probabilistic generation of various types of committed progenitors, each predetermined to give rise to an invariant set of descendant cell types (**Figure 1B**). In such a program, each type of progenitor would produce a characteristic distribution of descendant cell fates, introducing cell fate correlations on lineage trees. Identifying these characteristic distributions—or lineage motifs—could allow inference of otherwise hidden progenitor states. Further, spatial variation in the frequency with which a given motif appears could provide a mechanism for indirect modulation of cell type frequencies across space (**Figure 1C**).

Previous studies of cell lineage have focused predominantly on clonal tracing, which identifies descendants of a single cell but does not resolve their full tree of lineage relationships. Recently, new methods have begun to allow more complete lineage tree reconstruction. Time-lapse imaging allows direct tracking of lineage trees of differentiating progenitors ^16^. In addition, a new generation of engineered lineage reconstruction systems has emerged, which use CRISPR or recombinases to progressively edit barcode, or ‘scratchpad,’ sequences integrated in the genome ^17–23^. These edits accumulate stochastically in each cell over multiple cell cycles. Readout of endpoint edit patterns in individual cells allows reconstruction of their lineage relationships in a manner analogous to phylogenetic reconstruction. As these methods grow in scale and temporal resolution, they provoke the question of how fully resolved lineage trees with endpoint cell fates can be used to infer underlying developmental programs.

To address this challenge, we introduce Lineage Motif Analysis (LMA), a computational approach for inferring statistically overrepresented patterns of cell fates on lineage trees. LMA is based on motif detection, which has been used to identify the building blocks of complex regulatory networks ^24^, DNA sequences ^25, 26^, and other biological features ^27, 28^, but has not to our knowledge been applied to understand developmental programs. As a ‘bottom-up’, data-driven approach, LMA does not require specific assumptions about underlying molecular mechanisms and can be applied to diverse systems for which sufficient cell lineage information is available. Biologically, motifs could be generated by progenitors intrinsically programmed to autonomously give rise to specific patterns of descendant cell fates. They could also reflect more complex developmental programs involving extrinsic cues and cell-cell signaling that generate correlated cell fate patterns on lineage trees.

Here, we first define LMA and demonstrate how accurately it performs using simulated datasets. We then identify lineage motifs in published zebrafish and rat retina development datasets, as well as a dataset of early mouse embryonic development. These results reveal temporal and spatial differences in cell fate determination across different progenitors. Further, the appearance of shared retinal motifs across different species suggests that motifs may be evolutionarily conserved features of development. Computationally, we demonstrate how the use of lineage motifs facilitates adaptive variation in retinal cell type composition and show that this theory is consistent with known variation in vertebrate retinal cell type proportions. Together, these results support LMA as a broadly useful tool to understand developmental programs.

**Figure 1:**
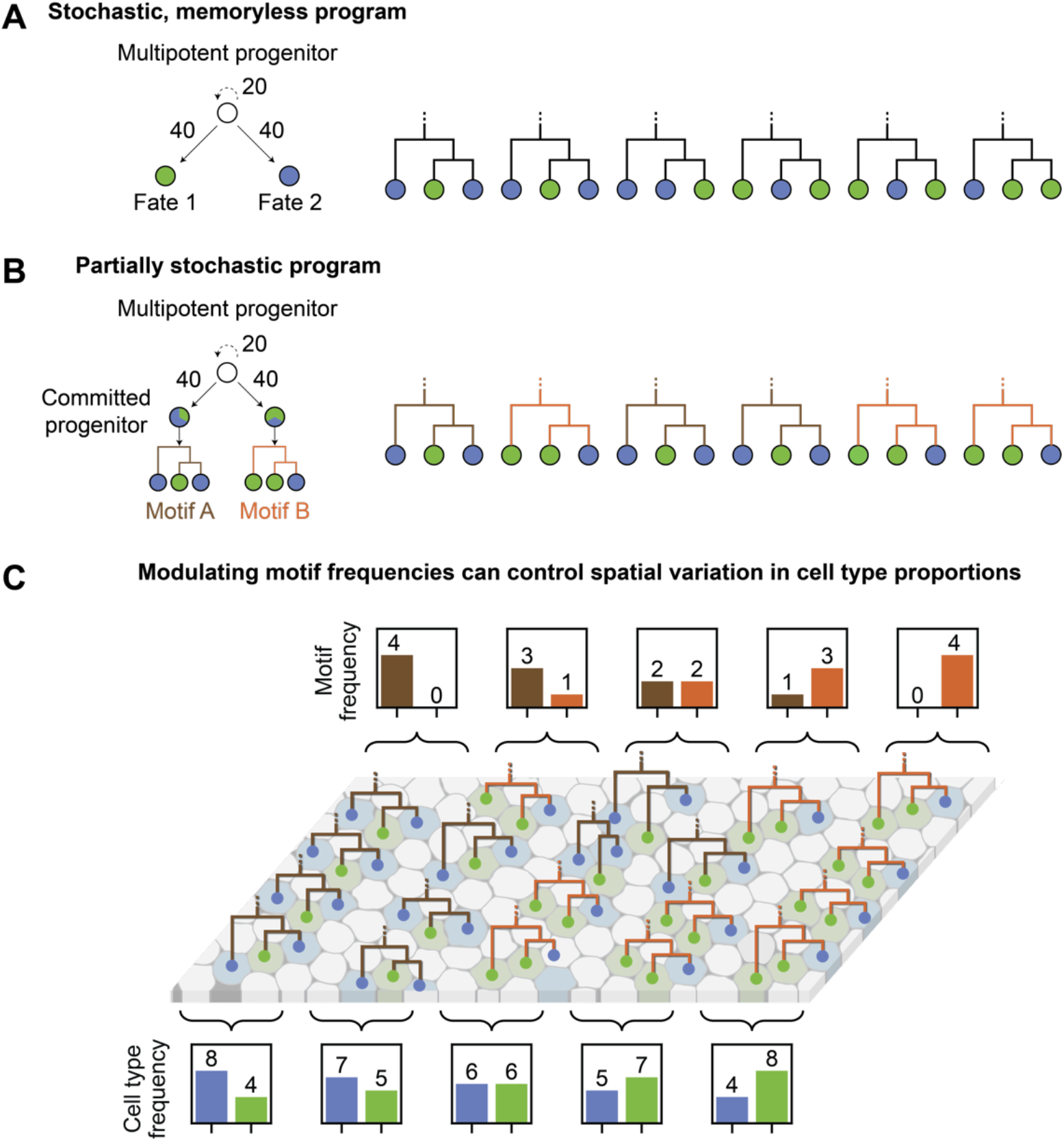
Cell type proportions can be controlled using partially stochastic programs that specify defined groups of cell types as motifs. A. An example of a completely stochastic developmental program where a multipotent progenitor can self-renew (with 20% probability) or give rise to different fates (with 40% probability each) in a memoryless manner. Lineage trees generated under this program would not exhibit fate correlations between related cells. B. An example of a partially stochastic program where a multipotent progenitor can self-renew (with 20% probability) or give rise to different types of committed progenitors (with 40% probability each). The committed progenitors differentiate to give rise to a defined set of cell types (motif A or B). Lineage trees generated under this program would exhibit fate correlations between related cells, representative of the committed progenitors present within the program. C. In this schematic, five spatial regions along the horizontal axis of a tissue are generated primarily through two types of triplet motifs (motif A or B). Depending on how frequently each motif is utilized, cell type proportions across the tissue can vary, but this variation is capped such that each cell type can at most be twice as abundant as the other type.

## Design

A previous study analyzed sister cell fate correlations by comparing the frequency of two-cell clones to that predicted by random association of cell types given their observed proportions ^15^. Another study analyzed triplet fate correlations by comparing the frequency of triplet patterns to that observed in simulated lineage trees using a stochastic model where each starting progenitor can self-renew or differentiate into all possible cell types within the dataset under a set of probabilities ^29^. These studies provide evidence for fate correlations between related cells. However, a framework that can be recursively applied to any lineage tree dataset to systematically identify lineage motifs of varying size remains lacking.

We first simulated sets of lineage trees with 3 different cell types (**Figure 2**). We enumerated all possible cell fate patterns, tabulated the number of times each occurred within the simulated trees, and compared these abundances to those expected in a “null” model. The null model was constructed by repeatedly resampling the cell fate labels on the simulated lineage trees (**Methods**). This procedure maintains overall cell fate proportions but eliminates fate correlations between related cells. To detect larger motifs, it is necessary to control not only for overall cell type frequencies but also for the frequencies of any “sub-patterns” within the pattern of interest. For example, a triplet pattern comprising a sister cell doublet and their common cousin could appear over-represented solely because the sister doublet is itself a motif. To account for this, we resampled in a manner that preserves sub-pattern frequency, by drawing from a pool of similar sub-patterns across all trees. For each pattern, we computed a z-score to quantify the degree of over-representation, as well as a false discovery rate adjusted p-value ^30, 31^ to measure significance. Finally, anti-motifs, defined as patterns that are underrepresented in the observed trees, were identified using the same approach.

LMA is distinct from a related approach termed kin correlation analysis (KCA). KCA infers cell state transition dynamics from lineage trees and endpoint cell state datasets, but is mainly applicable to systems governed by Markovian dynamics, in which sister cell transitions are independent of one another ^32, 33^. LMA also contrasts with the “lineage complexity” metric, which enumerates the minimal set of patterns necessary to describe the overall developmental program, but does not quantify how statistically overrepresented each pattern is within the dataset ^34^.

**Figure 2:**
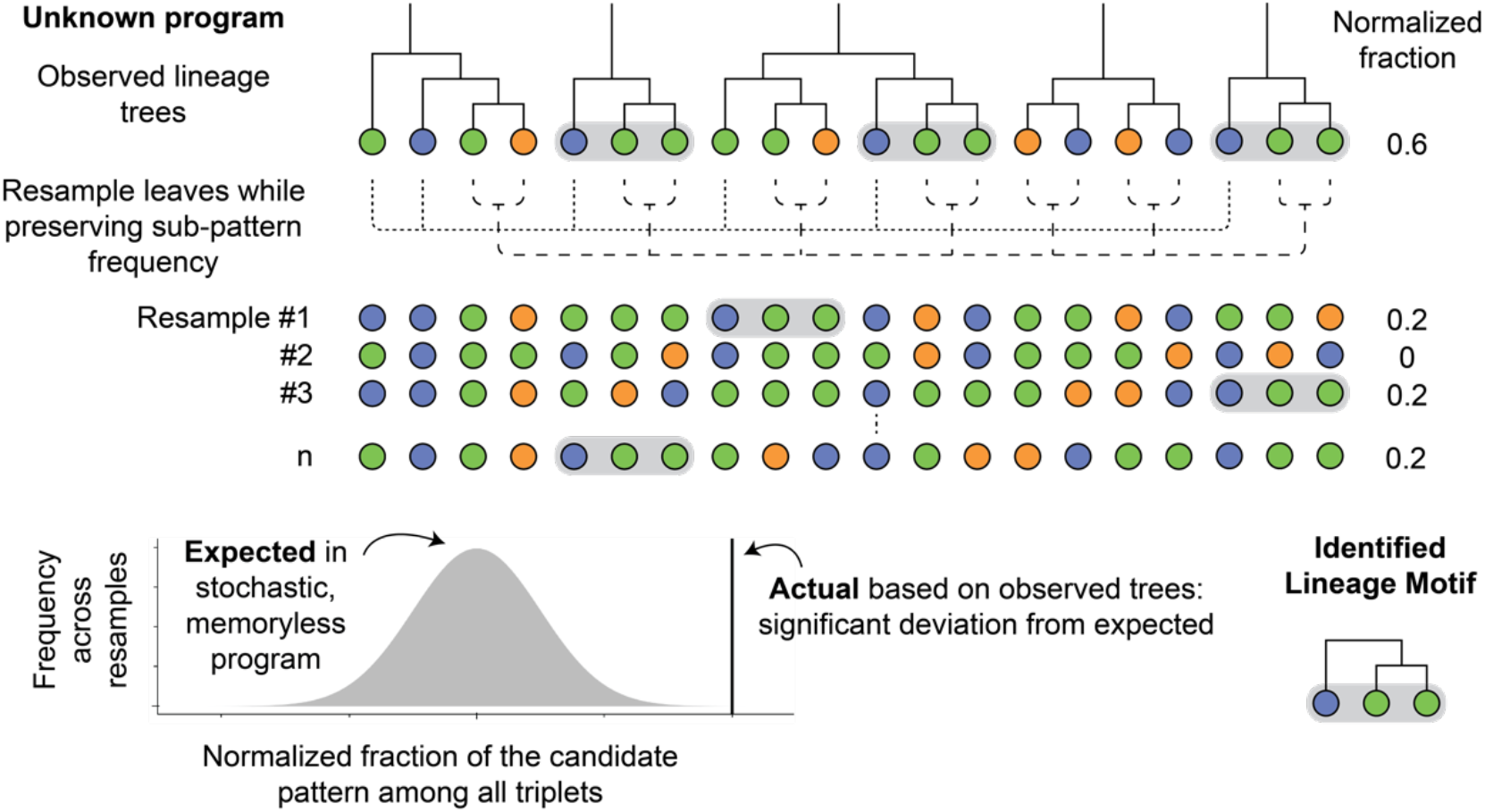
Lineage Motif Analysis (LMA) identifies fate correlations in lineage trees by statistical resampling. Statistically overrepresented patterns of cell fates on lineage trees can be identified by comparing the observed set of trees to those that have been resampled to eliminate fate correlations while maintaining sub-pattern frequency. An example of triplet lineage motif identification is shown here, where singlets and doublets are resampled across all trees. Candidate patterns that display significant deviation from the expected frequency based on the resampled trees are identified as lineage motifs.

To demonstrate that LMA can recover different types of committed progenitors present in larger developmental programs, we simulated lineage tree datasets using either a competence progression program (**Figure 3A** **and Figure S1**), reminiscent of neural developmental systems like retina, or a binary fate program (**Figure S2A and S3**), reminiscent of early embryonic development. For these programs, we used differentiation probabilities that generate roughly equal cell fate proportions in the overall dataset (**Figure 3B** **and Figure S2B**). We then applied LMA to the simulated tree datasets. In both cases, the resulting motifs reflected the structure of the underlying program and captured multiple levels of progenitor commitment over time. For example, in trees generated using a competence progression model (**Figure S1**), where cell fates A through F are generated progressively over time, only symmetric doublet patterns, such as (F,F) and (E,E), were statistically overrepresented within all possible doublet patterns (**Figure 3C**).

We next analyzed triplet patterns, in which a single progenitor divides to produce a terminal cell, X, and a second progenitor cell that divides once more to produce a doublet of terminal cells, Y and Z, producing a triplet denoted as (X,(Y,Z)). After accounting for both singlet and doublet frequencies, only triplet patterns including two sequential levels of progenitor commitment, such as (E,(F,F)) and (D,(E,E)), were significantly overrepresented (**Figure 3D**).

LMA can be scaled up to analyze larger asymmetric patterns, including quartets, quintets, sextets, and septets, which respectively span 3, 4, 5, and 6 cell divisions. Given a reasonable number of trees (500 total), the motifs successfully captured up to 5 levels of the competence progression program. Like the triplet results, the significant higher order motifs exclusively involved consecutive cell fate patterns. For example, (D,(E,(F,F))) was a motif, while (C,(E,(F,F))) was not. As motif size grows larger, the size of the dataset required for detection also increases (**Figure 3E**). Together, these results confirm that LMA can be used to recursively identify lineage motifs in large patterns.

We also analyzed trees generated using a binary fate model in which progenitors make binary choices which restrict their fate potential over time (**Figure S2A and S3**). The doublet and quartet motifs reflect the structure of the underlying program as expected (**Figure S2C-D**). However, no octet patterns were significantly over- or under-represented for the indicated rates of differentiation and self-renewal (using datasets up to 50000 trees; **Figure S2E-F**). Taken together, these results indicate that LMA is capable of recursively identifying lineage motifs of multiple sizes in different models of development and is especially powerful when applied to the competence progression dynamics, likely due to the lower number of possible patterns per level of progenitor commitment.

Finally, to enable the identification of lineage motifs across diverse developmental contexts, we created an open-source Python package, termed “linmo,” for identifying motifs in lineage trees. The package is available on a GitHub repository (https://github.com/tranmartin45/linmo), which includes supporting documentation and tutorials for processing the following lineage tree datasets analyzed here.

**Figure 3:**
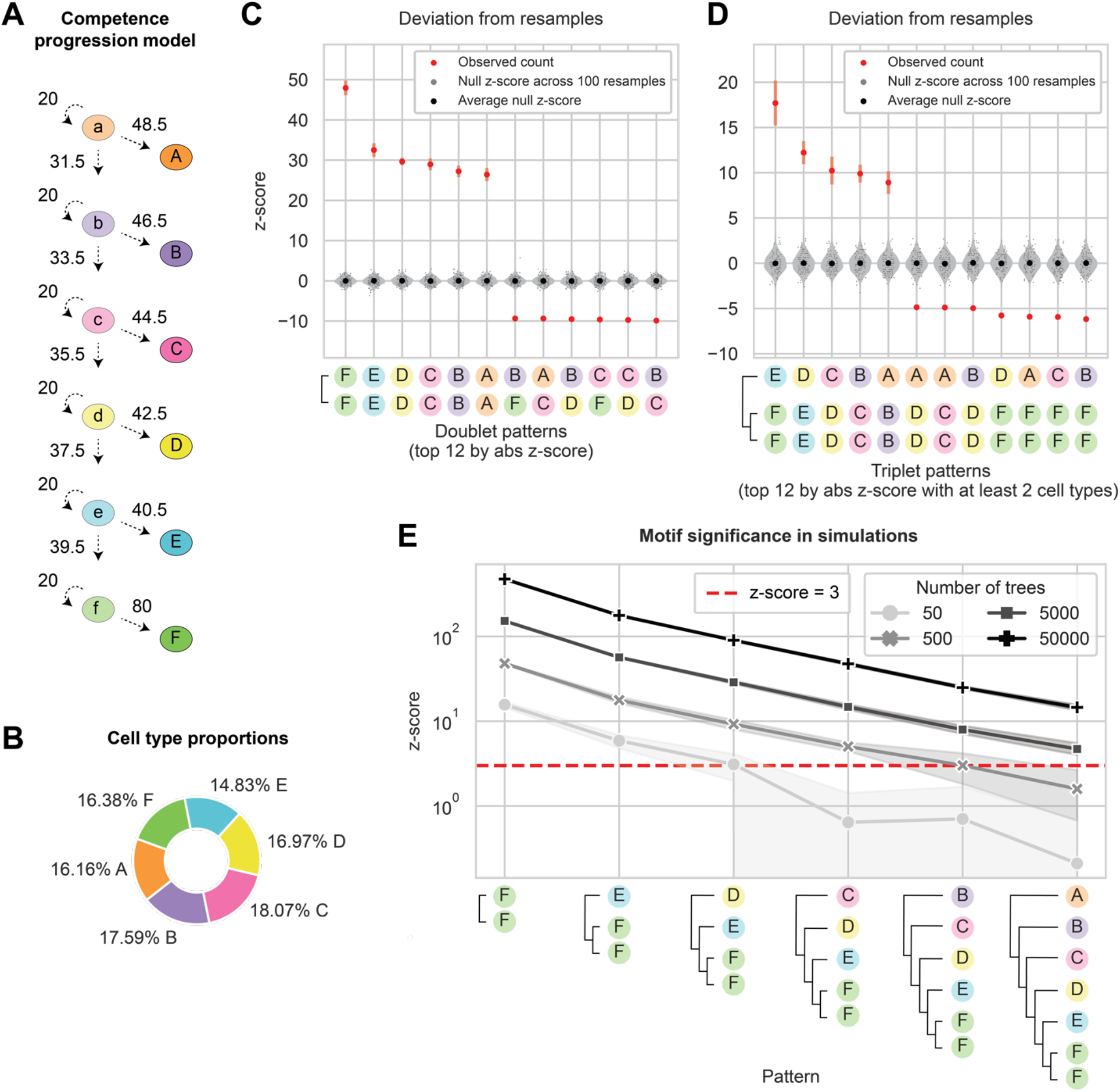
Motifs reveal committed progenitors in a competence progression model of development. A. Lineage trees were simulated using a competence progression model of development. Each progenitor can either self-renew with 20% probability, differentiate into a terminal fate, or progress to the next competence state (except for the last progenitor ‘f’). Probabilities for differentiating into a terminal fate or progression to sequential competence states were chosen to generate roughly equal cell type proportions. B. Cell type proportions in 500 simulated lineage trees. C. Deviation score for top 12 most significant doublet patterns, calculated using the mean and standard deviation of counts across 10000 resamples. Null z-scores were calculated by comparing a random resample dataset to the rest of the resample datasets. 10 datasets containing 500 simulated trees each were used, with the standard deviation across the datasets plotted as error bars. D. Deviation score for top 12 most significant triplet patterns with at least 2 cell types. E. Deviation score for select patterns that reflect sequential differentiation of cell fates using datasets of varying size. Shading indicates 95% confidence interval across 10 datasets for each point. See also **Figure S1-3**.

## Results

### LMA reveals spatial organization of fate commitment in zebrafish retina development

Retina development provides a well-studied example of cell fate diversification. It involves generation of a conserved set of terminal cell fates across diverse vertebrate species. At the same time, it also exhibits substantial inter-species variation in the spatial organization of cell types ^1^, making it an ideal target tissue for LMA. Therefore, we examined a zebrafish retina development dataset spanning 32 to 72 hours post fertilization (hpf) ^35^, during which progenitors terminally differentiate to form the major neuronal and glial cell types, including ganglion (G), amacrine (A), bipolar (B), photoreceptor (R), horizontal (H), and Müller glia (M) (**Figure 4A**). He et al. used time-lapse confocal microscopy in reporter zebrafish lines to track every cell division event for 60 retinal progenitors spanning the nasal-temporal axis. Their data supported previous work showing that a wave of differentiation starts in the nasal region and gradually progresses to the temporal region ^36, 37^. The cell type composition within clones was generally observed to be variable, with weak fate correlations between related cells. A key exception, however, was the frequent appearance of symmetric terminal pairs of photoreceptor, bipolar, and horizontal cells.

We partitioned lineage trees based on the progenitor spatial location and applied LMA, beginning with doublet patterns. We found that the (H,H), (B,B), and (R,R) doublet motifs were overrepresented in a similar manner across the three different retinal regions (temporal, middle, and nasal) (**Figure 4B-E**). The key exception was a lack of (H,H) and (B,B) doublets in the nasal region, likely because those cell types were only present at very low levels in this region (**Figure 4D-E**). These results were consistent with key findings from He et al., while extending the analysis to assess regional variation.

LMA also found new motifs not previously identified in the He et al. study and revealed how their frequency varies across space. For example, even though bipolar and amacrine cells appear at similar frequencies across all three retinal regions, the (A,B) doublet was specifically overrepresented in the nasal region (**Figure 4E**). Also, doublets comprising one R cell and all other cell types were generally underrepresented across all regions, constituting anti-motifs. We also searched for higher order motifs but found that no patterns were significantly over- or under-represented, indicating that higher order motifs are either absent or require larger datasets for detection (**Figure S4**). Overall, the observed motif profile suggests that amacrine and bipolar cells frequently share a common progenitor at the terminal cell division, specifically in the nasal region of the zebrafish retina, whereas photoreceptor and non-photoreceptor cells do not share a common progenitor at the terminal cell division in all regions.

**Figure 4:**
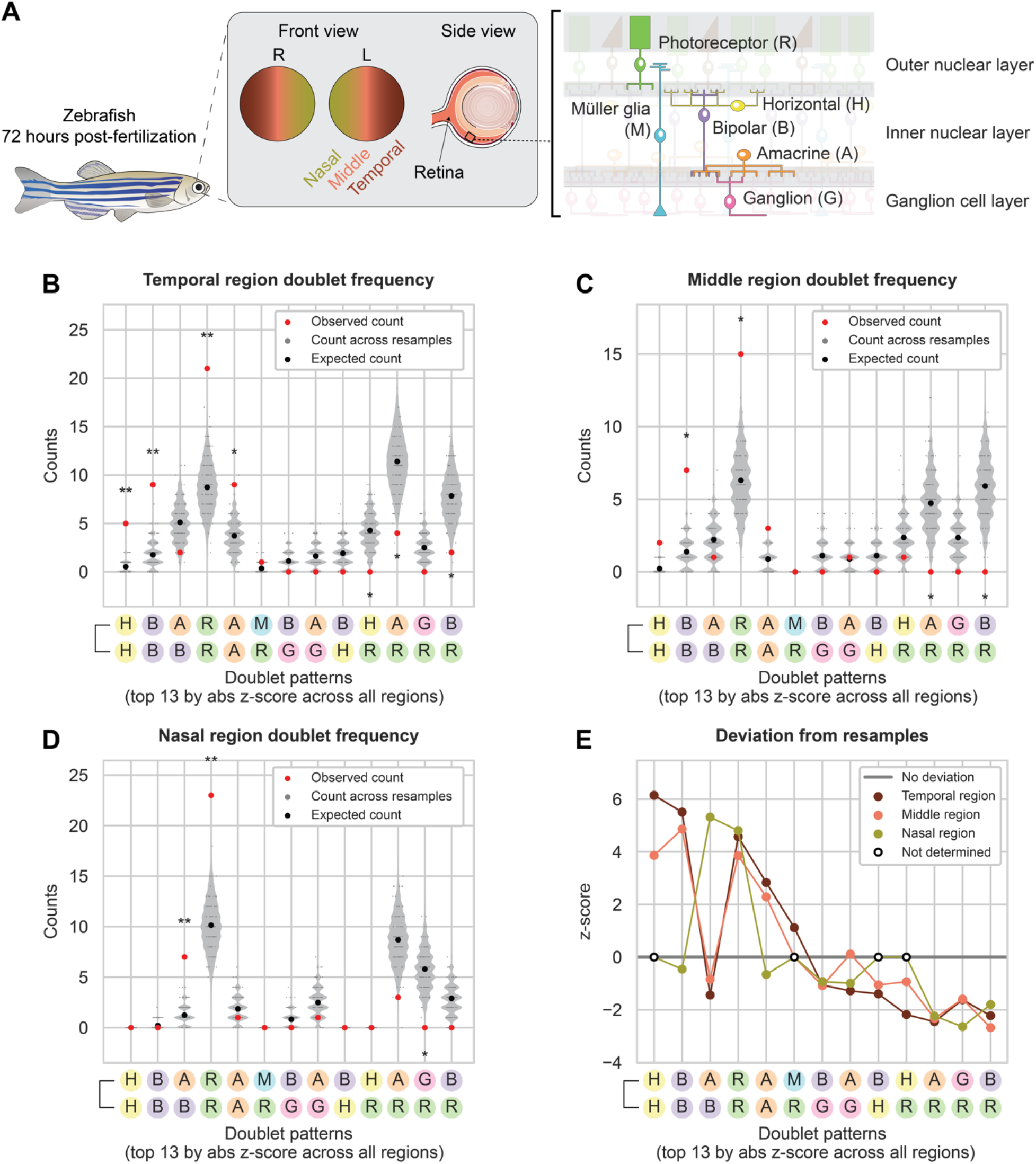
Doublet lineage motif analysis in zebrafish retina development shows spatial organization of fate commitment. A. The zebrafish retina contains five major types of neurons and Müller glia organized into three cell layers. Differentiation starts from the nasal side of the eye and progresses to the temporal side. B. Counts for doublet patterns in the observed zebrafish retina trees from He et al. ^35^ in the temporal region and across 10000 resamples (* = adjusted p-value < 0.05; ** = adjusted p-value < 0.005). All 10000 resamples are represented in the violin plots, but a random subset of only 100 resamples are shown as overlaying dot plots. The top 13 significant doublets across progenitors from all spatial regions are shown. The expected count was calculated analytically (**Methods**). C. Counts for doublet patterns in the middle region of zebrafish retina and across 10000 resamples. D. Counts for doublet patterns in the nasal region of zebrafish retina and across 10000 resamples. E. Deviation score for doublet patterns in the temporal, middle, and nasal region, calculated using the mean and standard deviation of counts across 10000 resamples. Doublet patterns with an observed and expected count of 0 were omitted from analysis. See also **Figure S4**.

### Shared retinal lineage motifs across species suggest conservation of developmental programs

Are retinal lineage motifs conserved between different species? To address this question, we analyzed a dataset of postnatal rat retinal progenitor cells grown *in vitro* at clonal density, consisting of 129 lineage trees with at least 3 cells ^29^. During this period, rat retinal progenitor cells gave rise to mostly rod cells (R, 74.6% of cells), some bipolar and amacrine cells (respectively B and A, with 12.6% and 10.1% of cells), and few Müller glia (M, 2.7% of cells) (**Figure 5A**). In this work, the authors showed that a stochastic model based on independent fate decisions could explain the observed frequencies of most triplet patterns, arguing against the existence of specific fate programs ^29^.

Applying LMA to this rat retina dataset confirmed some of these conclusions, such as over-representation of (B,(A,B)) triplets (**Figure 5C, E**). However, it also revealed additional features of rat retinal development. For example, using LMA, we found that (A,B), (B,M), and (A,A) doublets were overrepresented while (B,R) doublets were underrepresented (**Figure 5B, D**). Correcting for sub-pattern frequencies in the triplet analysis revealed that the apparent over-representation of the (R,(A,A)) triplet in the previous study ^29^ could be entirely explained by the (A,A) doublet motif frequency. This highlights the importance of the recursive nature of LMA (**Figure 2, Methods**). Furthermore, because the progenitors were grown at *in vitro* at clonal density to minimize the effect of extrinsic cues on fate commitment, these motifs likely represent intrinsic developmental programs that generate predetermined sets of cell fates on lineage trees.

We next compared the motif profile between zebrafish and rat retina. We limited this analysis specifically to cell types that are shared between the analyzed datasets. Notably, the (A,B) and (A,A) motifs and the (B,R) anti-motif are observed in both species, suggesting that developmental programs are evolutionarily conserved. In contrast, the (B,B) and (R,R) motifs appear specifically in the zebrafish retina, while the (B,M) motif appears specifically in the rat retina. Overall, these data suggest that cell fate allocation in retina across species can occur in a deterministic and evolutionarily conserved manner, in which amacrine and bipolar cells share a common progenitor at the terminal cell division. At the same time, other programs may be more species-specific. For example, bipolar and Müller glia tend to share a common progenitor in rat, but not zebrafish, retina at the terminal cell division. More generally, these results provide a case example for how LMA can be used to assess the evolution of developmental programs.

**Figure 5:**
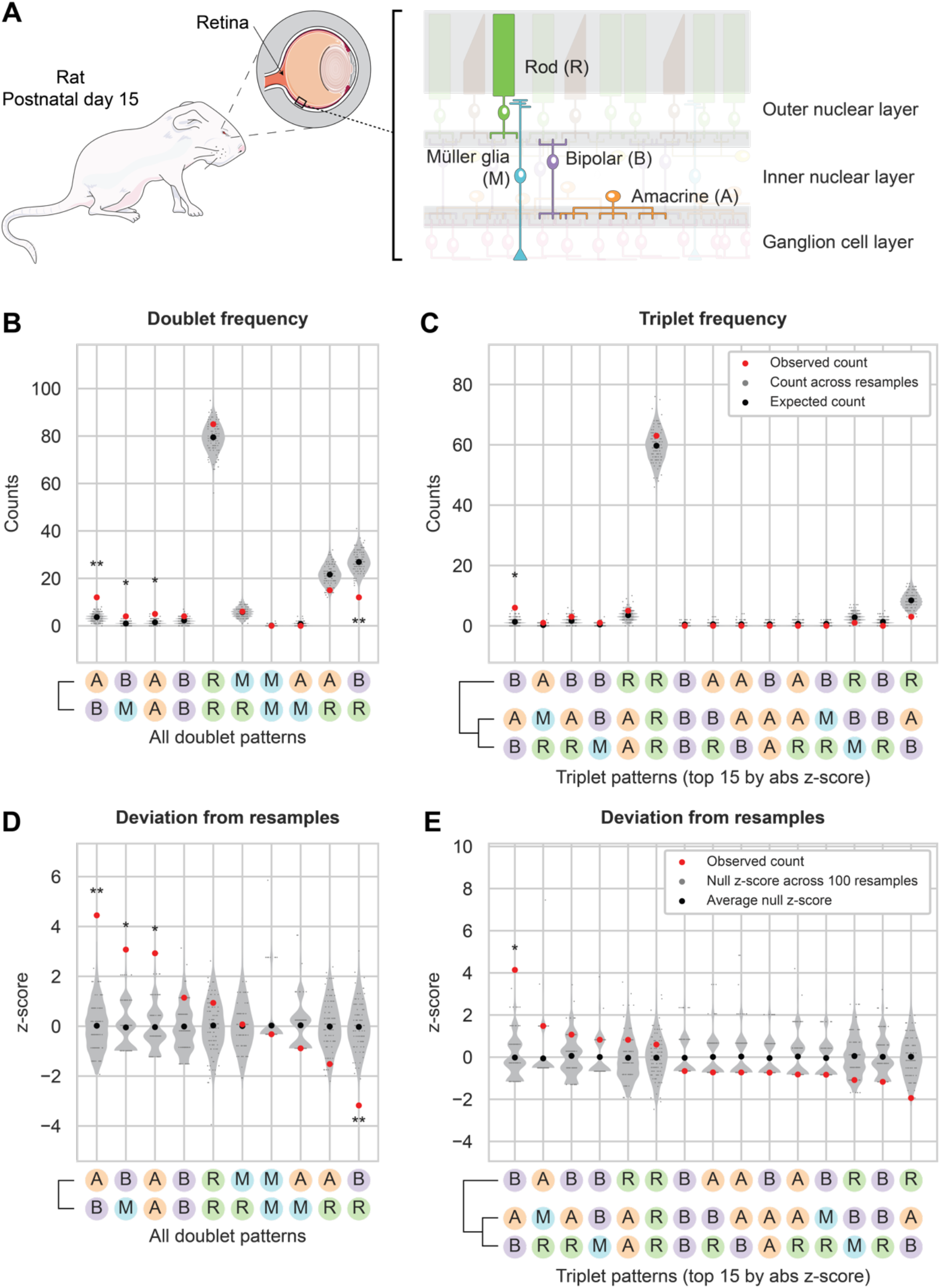
Doublet and triplet lineage motif analysis reveals fate commitment patterns in rat retina development. A. Schematic shows the cellular architecture of the mammalian retina. Cell types used in this study are highlighted. B. Counts for all doublet patterns in the observed lineage trees from Gomes et al. ^29^ and across 10000 resamples (* = adjusted p-value < 0.05; ** = adjusted p-value < 0.005). All 10000 resamples are represented in the violin plots, but a random subset of only 100 resamples are shown as overlaying dot plots. The expected count was calculated analytically (**Methods**). C. Counts for the top 15 significant triplet patterns in the observed lineage trees and across 10000 resamples. D. Deviation score for all doublet patterns in the observed lineage trees, calculated using the mean and standard deviation of counts across 10000 resamples. Null z-scores were calculated by comparing a random resample dataset to the rest of the resample datasets. E. Deviation score for the top 15 significant triplet patterns in the observed lineage trees.

### LMA reveals temporal differences in fate commitment during early mouse embryo development

Early embryonic development features conserved cell types across mammals and spatially restricted cell fate specification, making it an ideal system to apply LMA. We therefore used LMA to analyze a dataset of early mouse embryo development, spanning the 8-cell stage to blastocyst ^38^. During this period, cells make two major cell fate decisions. The first fate decision distinguishes between inner cell mass (ICM) and trophectoderm (T). Subsequently, ICM cells either undergo apoptosis (A) or further differentiate into either epiblast (E) or primitive endoderm (P) fates (**Figure 6A**).

In a previous study, Morris et al. ^38^ used time-lapse confocal microscopy to trace individual progenitor cells starting at the 8 to 16-cell division within 20 mouse blastocysts until their final fate is known at the late blastocyst stage (∼E4.5). The authors found that progenitors that internalize during the 8-16 cell stage are biased to give rise to epiblast (E), whereas those that internalize during or after the 16-32 cell stage are biased to give rise to primitive endoderm (P). However, it remained unclear whether individual progenitors give rise to sets of correlated cell fates in the final few divisions prior to the late blastocyst stage.

To gain insight into this question, we partitioned lineage trees into those generated from inside or outside progenitors at the 16-cell stage and applied LMA to both sets. Examining the final cell division, we found that most doublet patterns (90%) within both types of progenitors are either motifs or anti-motifs (**Figure S5**). For example, symmetric sister pairs such as (P,P), (E,E), and the apoptotic doublet (A,A) were over-represented among descendants of both inside and outside progenitors, and therefore motifs. (T,T) was also a motif among outside progenitors (inside progenitors do not give rise to trophectoderm). These results suggest that by E4.5, most (76.5%), but not all, cells have already committed to one of the three lineages before the previous cell division, and therefore produce symmetric doublets.

We also observed asymmetric doublet motifs, such as (A,P), (A,E), (E,P), which were overrepresented in trees from outside progenitors while underrepresented in trees from inside progenitors. Trophectoderm, unlike the other lineages, was part of asymmetric anti-motifs among outside progenitors. Overall, the weaker motif signatures for inside progenitors suggest less commitment compared to the strong, and usually symmetric, doublet motifs among the descendants of the outside progenitor cells.

The recursive nature of LMA allowed us to extend it to identify higher order motifs, starting with triplets, in these data (**Figure 6**). Strikingly, we observed triplet motifs with multiple cell fates, such as (A,(P,P)) and (T,(A,P)). In order to qualify as a motif, these patterns must occur more often than expected after accounting for the frequencies of their sub-patterns. To achieve that level of over-representation suggests the existence of progenitors that are biased to produce complex three-cell patterns. Biologically, the overrepresented (T,(A,P)) triplet could represent a committed outside progenitor at the 16-cell stage, which divides asymmetrically to give rise to a trophectoderm cell and an internalized intermediate progenitor, which then gives rise to an apoptotic and primitive endoderm cell. Among outside cells, we also observed homogeneous triplet motifs, including (P,(P,P)) and (E,(E,E)), suggesting the existence of fully committed progenitors at least two generations earlier. We sought to detect quartet motifs (**Figure S6**), but the dataset was too small to reliably detect significant deviations from null expectations. Taken together, these results suggest that some outside progenitors may commit to give rise to defined groups of cell types at least two cell divisions before the late blastocyst stage, while inside progenitors remain plastic and uncommitted towards certain fates.

**Figure 6:**
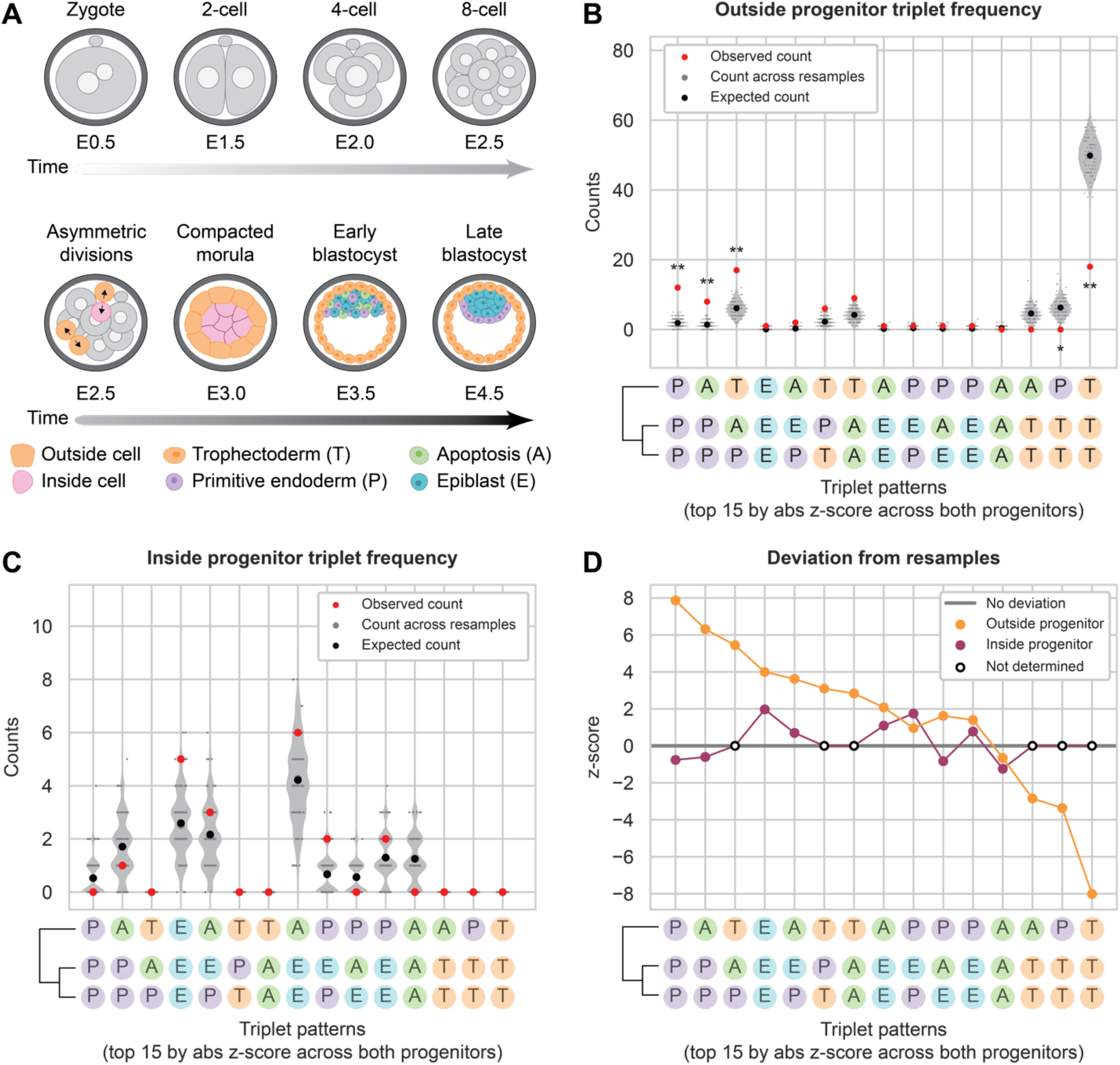
Triplet lineage motif analysis in mouse blastocyst development suggests outside progenitors initiate fate commitment earlier than inside progenitors. A. Early in mouse development, blastomeres are partitioned into outside and inside progenitors based on their spatial position in the compacted morula. As development progresses, outside cells form trophectoderm (T), while inside cells either undergo apoptosis (A), or contribute to epiblast (E) or primitive endoderm (P). B. Counts for triplet patterns in the observed mouse blastocyst trees from Morris et al. ^38^ in the outside progenitors and across 10000 resamples (* = adjusted p-value < 0.05; ** = adjusted p-value < 0.005). All 10000 resamples are represented in the violin plots, but a random subset of only 100 resamples are shown as overlaying dot plots. The top 15 significant triplets across both sets of progenitors are shown. The expected count was calculated analytically (**Methods**). C. Counts for triplet patterns in the observed mouse blastocyst trees in the inside progenitors and across 10000 resamples. D. Deviation score for triplet patterns in the outside and inside progenitors, calculated using the mean and standard deviation of counts across 10000 resamples. Triplet patterns with an observed and expected count of 0 were omitted from analysis. See also **Figure S5-6**.

### Lineage motifs facilitate adaptive variation in cell type frequencies

How do biological systems facilitate adaptive variation in cell type frequencies, either spatially within a tissue, or evolutionarily between species? Within the large space of potential cell type distributions, most are likely to be maladaptive. Is it possible to structure the developmental program in such a way that allows cell type frequencies to mainly vary within adaptive, or optimal, regimes ^3, 4^?

We reasoned that lineage motifs could address this problem. Mathematically, lineage motifs represent a linear transformation from a set of motif frequencies to a set of cell type frequencies. Each motif generates a subset of cell types in a particular stoichiometric ratio. For example, the (A,B) doublet motif generates bipolar and amacrine cells in a 1:1 ratio (**Figure 5D**). If most cell fate decisions were controlled through motifs, then a developing tissue could indirectly control the frequencies of individual cell types by specifying the frequency of each motif (**Figure 1C**).

More precisely, we can describe the conversion from motif frequencies to cell type distributions as a linear transformation: *z*(*s*) = *X* ∗ *y*(*s*) + *e*(*s*). Here, *z*(*s*) is a vector whose components represent the counts of each cell type in position/species *s*, *X* is a non-negative integer matrix describing how many cells of each type (rows) are produced by each motif (columns), *y*(*s*) denotes the motif frequencies in position/species *s*, and *e*(*s*) represents the number of additional cells of each type in position/species *s* that cannot be explained through the motifs (**Figure 7A**). In defining *X*, it is important to note that certain fate patterns may be overrepresented, and be detected as motifs, in one spatial region or species but not another. A complete set of motifs, and therefore a complete specification of *X*, would include motifs identified in all biological contexts.

To understand how motifs constrain cell type frequencies, we first constructed a set of hypothetical motif matrices *X*_0_, *X*_1_, *X*_2_, each reflecting a different motif structure. *X*_1_ is a diagonal matrix representing the null model in which each column trivially corresponds to a single cell type. In contrast, *X*_1_ and *X*_2_ respectively contain exclusively doublet or triplet motifs where multiple cell types are generated together (**Figure 7B**). For each motif matrix, we simulated datasets by randomly choosing frequencies for each of the motifs present within each matrix (**Methods**). For simplicity, we initially assumed *e* = 0 and constrained cell type frequencies to sum to a constant, ∑_i_ *z*_i_ = *const.*, to reflect the limited total capacity of the tissue. We then analyzed the range of cell type distributions produced by each matrix. Under the null model, cell type frequencies spanned the full space, as expected. By contrast, the other two models restricted cell type frequencies to limited subspaces. This can occur, for example, by excluding distributions consisting mainly of only one cell type.

Although motifs can constrain cell type distributions in general, it remained unclear whether the specific motifs observed in the rat retina dataset would be consistent with the distributions of retinal cell types independently observed in different species. Addressing this question requires (1) defining the motif-accessible space of cell type frequencies permitted by the observed rat retina motifs, and (2) determining whether independently measured retinal cell type proportions from other vertebrate species lie within that space.

To define the motif-accessible frequency space, we first need to determine the lower and upper bounds for cell type frequencies. We set the lower bounds at *e*_3_, the cell type counts for all cells in the rat retina dataset born outside of a motif **(Methods, Figure S7A)**. We set the upper bounds by constraining the total number of cells to be the same as in the rat retina dataset, ∑_i_(*z*_3_)_i_ = *const*. Using these constraints, we simulated datasets containing randomly chosen frequencies for each of the observed rat retina motifs in *X*_3_, or as a control, the null model, *X*_0_. The motif model accessed only a subset of the space of fate proportions spanned by the null model (**Figure 7C**). Within this subspace, the motif model showed higher density of fate distributions corresponding to moderate levels of both amacrine and bipolar cells and low levels of Müller glia. Bipolar cells and Müller glia exhibited a reduced maximum proportion relative to the null model, consistent with the observation that both cell types are generated with other cell types in the rat retina motifs.

We compared the datasets generated using the motif or null model to independent measurements of retinal cell type proportions across multiple vertebrate species ^2, 39^. Strikingly, this analysis revealed that all the independently measured fate distributions of vertebrate retina lie within, or very close to, the subspace accessed by the motif model. To understand how the structure of the motif matrix impacts the resulting cell type distributions, we repeated this analysis omitting the (A,A) motif from the motif matrix *X*_%_ (**Figure S7B**). This resulted in a smaller subspace achieved by the motif model, specifically lowering the maximum proportion of A cells from 62.8% to 46.0% (**Figure S7C**). This perturbed model failed to capture the empirical cell fate distributions for mouse, rabbit, monkey, and chick retina, indicating that the (A,A) motif is required to explain variation in cell type proportions across vertebrate retina.

Taken together, these results are consistent with the notion that motifs identified in lineage trees of rat retina could facilitate evolutionary variation in retinal cell type proportions across vertebrates. They further show that the range of cell type proportion space achieved using the motif model can be expanded or constrained by respectively increasing or decreasing the number of different motifs in the model. In the future, a more complete identification of motifs using larger datasets could therefore expand and modify the accessible space of cell type proportions described here.

**Figure 7:**
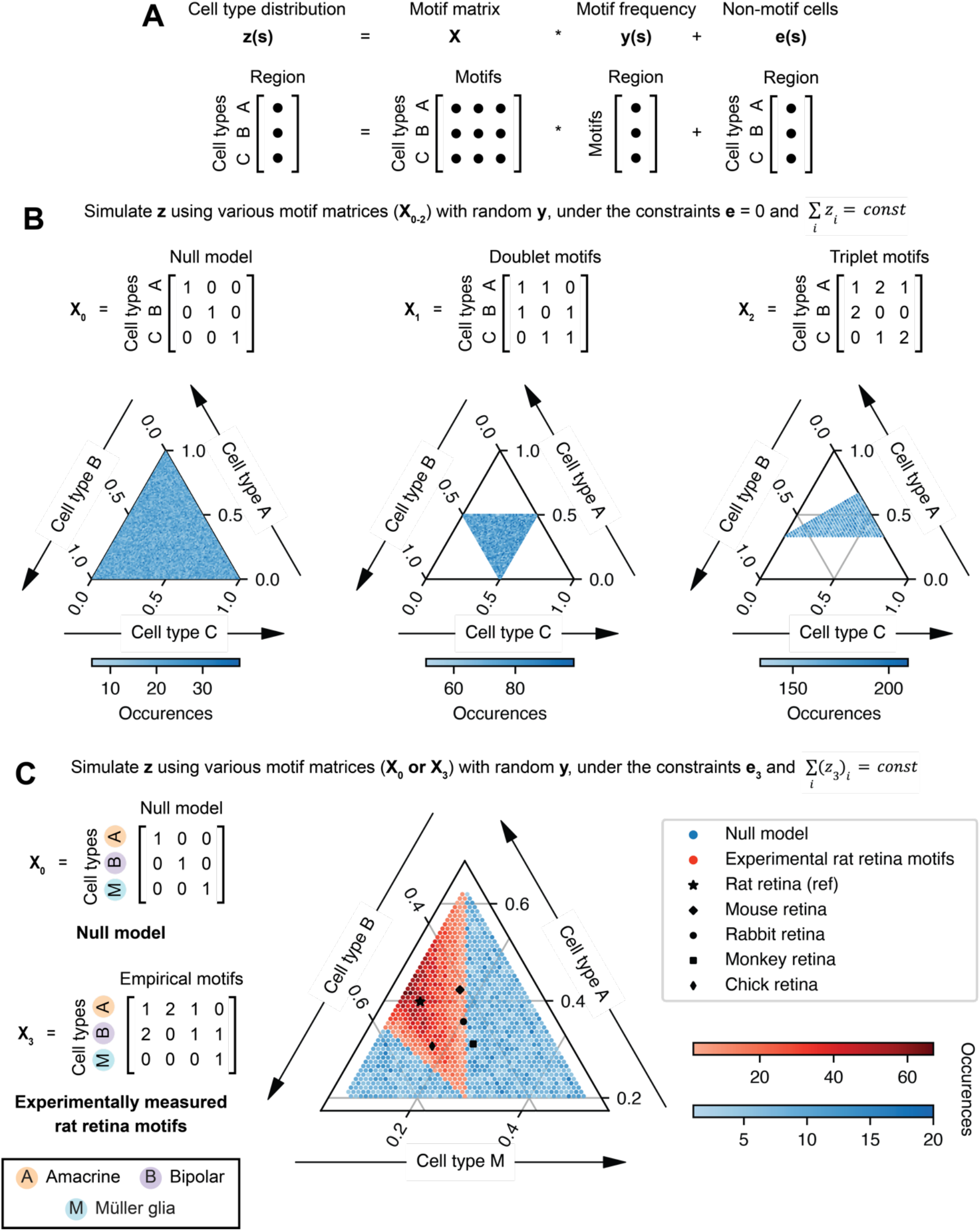
Motifs can facilitate optimal variation in cell type frequencies between species. A. The *z*(*s*) = *X* ∗ *y*(*s*) + *e*(*s*) matrix equation describes the linear transformation from motif frequencies to cell type distributions. B. Cell type distributions were simulated by randomly varying the frequencies of motifs using three example motif matrices (*X*_0_, *X*_1_, *X*_2_), assuming no cells are born outside of a motif (*e* = 0) and the total cell type frequencies to sum to a constant, ∑_i_ *z*_i_ = *const*. *X*_0_ corresponds to the null model, *X*_2_ corresponds to doublet motifs, and *X*_#_ corresponds to triplet motifs. The data was plotted as a ternary plot where each axis corresponds to the proportion of one cell type. C. Cell type distributions were simulated using a null model (*X*_0_) or the empirical motif matrix based on the rat retina motifs (*X*_3_) in Figure 5. The lower bounds were set at *e*_%_, the counts for all cell types in the experimental rat retina dataset that were born outside of a motif. The upper bound was set by constraining the total number of cells to be the same as in the rat retina dataset, ∑_i_(*z*_3_)_i_ = *const*. The data was plotted as a ternary plot where each axis corresponds to the proportion of one cell type, with the cell type proportions of mouse, rabbit, monkey, and chick retina from Masland ^2^ and Yamagata et al. ^39^ overlaid. See also **Figure S7**.

## Discussion

Producing cell types in optimal ratios is essential for tissue function. In many contexts, these proportions are established during development, when progenitor cells self-renew or differentiate into more committed progenitors or terminal fates. These committed progenitors can be identified through the repertoire of descendant cell types they produce. Increasing recent attention to the role of lineage in development ^40^ and the emergence of new methods for reconstructing lineage trees ^41, 42^ provoke the question of how one can infer developmental programs and different types of committed progenitors based on the arrangement of descendent cell fates on lineage trees.

In this work, we introduce a general computational approach, Lineage Motif Analysis (LMA), based on statistical resampling of lineage trees. Using simulations, we demonstrated that LMA can be recursively applied to uncover fate correlations in large patterns that span multiple cell divisions. By applying this framework to three biological datasets, we validated known fate patterns and identified novel fate correlations. In the retina, motifs can recur across space, or appear specifically in certain regions of the tissue. The presence of shared motifs across zebrafish and rat retina suggests evolutionary conservation of retina developmental programs. In the mouse blastocyst, inside progenitors appear plastic and less committed towards certain fates compared to outside progenitors during the final cell division before the late blastocyst stage. Based on triplet motifs, some outside progenitors at the 16-cell stage appear to have already committed towards defined groups of fates at least two cell divisions before the late blastocyst stage. Finally, we showed that the motifs identified in the rat retina dataset, if utilized in different proportions, could explain variation in cell type frequencies across several vertebrate species. Lineage motifs thus provide a useful and biologically meaningful lens through which we can analyze developmental programs.

Lineage motifs could be regarded simply as the consequence of a differentiation process that requires the cells to pass through intermediate states of partial fate commitment. However, this explanation still leaves open the question of why certain commitment states have been selected and why certain cell types appear in multiple motifs. One potential answer is that lineage motifs play functional roles in controlling cell type proportions. Because lineage motifs generate groups of cell types in fixed stoichiometric ratios, this could allow the organism to use motifs as ‘knobs’ that can modulate cell type proportions, while maintaining them within physiologically adaptive regimes. At the same time, we emphasize that developmental systems can use a multitude of other mechanisms for establishing and modulating cell type proportions, including morphogens, programmed cell death, quorum sensing, migration, etc. In the future, we anticipate greater availability of high-quality lineage datasets, which should allow more complete tabulation of motifs across different tissue contexts. These data should thus enable more stringent tests of the model proposed here.

A second potential role for lineage motifs could be to create spatially localized neighborhoods of interacting cell types to implement specific functions. In the context of the retina, particular types of interneurons must be synaptically connected. For example, in crossover inhibition, OFF bipolar cells receive input from ON amacrine cells, which are depolarized by ON bipolar cells at light onset^43^. A neural circuit of these cell types in close spatial proximity could be ensured by regulating the generation of these cell types through a lineage motif, such as the (B,(A,B)) motif observed in the rat retina dataset (**Figure 5E**). Consistent with this hypothesis, recent work has shown that specific synapses develop preferentially among sister excitatory neurons in the mouse neocortex ^44^. In future studies, it will be crucial to characterize the functions of individual cells within a lineage motif.

Lineage motifs can be compared with other methods for inferring developmental programs, such as pseudotime, where single cells are densely profiled throughout time to obtain a population-level branching continuum of cell states ^41^. A previous study involving pseudotime inference suggested that molecularly defined subpopulations of retinal progenitors give rise to different sets of cell types ^45^. In particular, neurogenic early stage progenitors give rise to ganglion, amacrine, and horizontal cells, Otx2+ late stage progenitors give rise to bipolar and rod cells, and other late stage progenitors give rise to Müller glia. However, in our analysis of both the zebrafish and rat retina, we observe progenitors that are biased to form a sister pair of one amacrine and one bipolar cell, i.e. the (A,B) doublet motif. In the rat retina, we also observe progenitors that are biased to form a sister pair of one bipolar cell and one Müller glia, i.e. the (B,M) doublet motif (**Figure 4 and 5**). Therefore, individual progenitors during development can generate lineage patterns that deviate from the population-level trajectories inferred using pseudotime ^46^.

Looking forward, LMA should be especially useful for contexts that have systematic spatial or cross-species variation in cell type composition, like the intestine, pancreas, and liver ^47–50^. Moreover, in diseased tissues where cell type proportions are misregulated, motifs may provide a way to infer underlying developmental mechanisms. Deeper tree reconstructions, enabled by recording systems with greater memory capacity ^41, 42^, could enable the analysis of lineage hyper-motifs, representing higher level correlations between constituent motifs^51^. Analyzing how signal or transcription factor dynamics are correlated with the generation of motifs will reveal how this process is extrinsically or intrinsically regulated during development. Overall, by decomposing complex developmental programs into their functional building blocks, lineage motifs should help provide new insights into longstanding questions in development and evolution.

### Limitations

Although we have identified motifs and anti-motifs in three different lineage tree datasets in this work, it is likely that not all underlying developmental programs will recur with high enough significance to be classified as a motif. Programs with weaker fate correlations and datasets of limited size can hinder motif identification. Additionally, incomplete identification of cell types due to the use of limited numbers of markers in the experimental studies analyzed here could prevent discovery of more complex motifs.

## Acknowledgements

We thank Duncan Chadly, Ben Emert, Jacob Parres-Gold, Yodai Takei, and other members of the Elowitz lab for scientific input and feedback. We thank Mykel Barrett, Marianne Bronner, Rusty Lansford, Carlos Lois, Magda Zernicka-Goetz, and Meng Zhu for helpful discussion and feedback. This research was supported by Paul G. Allen Frontiers Group and Prime Awarding Agency under Award No. UWSC10142 and the National Institute of Health grant R01MH116508. M.T. was supported with funding from the Tianqiao and Chrissy Chen Institute for Neuroscience at Caltech and by the National Eye Institute of the National Institutes of Health under Award Number F31EY033220. A.A. was supported with funding from National Eye Institute of NIH under Award Number R00EY031782. M.B.E. is a Howard Hughes Medical Institute investigator. The content is solely the responsibility of the authors and does not necessarily represent the official views of the NIH. This article is subject to HHMI’s Open Access to Publications policy. HHMI lab heads have previously granted a nonexclusive CC BY 4.0 license to the public and a sublicensable license to HHMI in their research articles. Pursuant to those licenses, the author-accepted manuscript of this article can be made freely available under a CC BY 4.0 license immediately upon publication.

## Author contributions

Conceptualization, M.T., A.A., and M.B.E.; Methodology, M.T., A.A., and M.B.E.; Software, M.T. and A.A.; Formal Analysis, M.T. and A.A.; Writing – Original Draft, M.T.; Writing – Review & Editing, M.T., A.A., and M.B.E.; Visualization, M.T.; Supervision, A.A. and M.B.E.; Funding Acquisition, M.T., A.A., and M.B.E.

## STAR Methods

### Resource availability

#### Lead contact

Further information and requests for resources should be directed to and will be fulfilled by the lead contact, Michael B. Elowitz.

#### Materials availability

This study did not generate new unique reagents.

#### Data and code availability

● This paper analyzes existing, publicly available data. These accession numbers for the datasets are listed in the key resources table.
● All code used in this study has been deposited at GitHub (https://github.com/tranmartin45/linmo) as well as the CaltechDATA research repository (https://doi.org/10.22002/htgfr-11t35) and is publicly available as of the date of publication. DOIs are listed in the key resources table.
● Any additional information required to reanalyze the data reported in this paper is available from the lead contact upon request.

### Method details

#### Lineage tree resampling and motif identification

NEWICK-formatted lineage trees were first sorted to have doublet and quartet patterns arranged in alphabetical order according to their cell type annotations. All patterns were then aligned in order of earliest to latest born cells. For example, before alignment, triplet patterns could be present in the raw lineage tree data as ((X,X),X) or (X,(X,X)), and were therefore aligned to match the latter format in all cases. A similar procedure was followed for higher-order patterns, like asymmetric quartets, quintets, sextets, and septets.

All cell types and cell type patterns were then enumerated and counted for the number of occurrences within the lineage trees. The datasets were then appropriately resampled according to the type of motifs to be identified. For doublet motif identification, each cell type in the lineage tree dataset was replaced by a random cell type drawing from a list of all cell types within the dataset. Our results were not sensitive to replacing with vs. without replacement. For triplet motif identification, each singlet and doublet in the lineage tree dataset was respectively replaced by a random singlet or doublet drawing from a list of all singlets or doublets in the dataset. In this way, the overall frequencies of singlets and doublets remains the same across the resampled dataset while eliminating fate correlations between particular singlets and doublets. For quartet motif identification, each doublet in the lineage tree dataset was replaced by a random doublet drawing from a list of all doublets in the dataset. A similar procedure was followed for increasingly larger patterns.

The occurrences of each pattern were counted for each resampled dataset, then used to calculate an average number of occurrences and standard deviation across all resamples. The average and standard deviation were then used to calculate a z-score as follows:

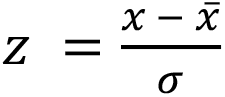

where *x* is the observed count in the original set of lineage trees, *x̄* is the average count across all resamples, and σ is the standard deviation across all resamples.

For plotting, the expected count of each pattern was calculated by multiplying the marginal probabilities of observing each of the two sub-patterns by the total number of that pattern across the entire dataset. Additionally, if the sub-patterns were not identical, the expected number was multiplied by two. For example, the expected number of the triplet (A,(B,C)) would be calculated as P(A) * P((B,C)) * 2, and the expected number of the quartet ((A,B),(A,B)) would be calculated as P((A,B)) * P((A,B)). The null z-scores were calculated by repeating the same resampling procedure above for randomly chosen resampled datasets.

#### Construction of synthetic lineage tree datasets

To test the recursive nature and accuracy of LMA in **Figure 3**, synthetic lineage tree datasets were simulated using a competence progression model or binary fate model of development. Each tree started as an ‘a’ or ‘i’ progenitor for each respective model, and a cell division was simulated producing two descendant cells, whose fates were chosen probabilistically based on the transition probabilities of the parental progenitor type. Cell divisions were repeatedly simulated for all progenitors present within the tree until all cells reached terminal fates (A-F or A-H for each respective model).

#### Simulation of cell type proportions with input motif matrices

Cell type proportions were first simulated using input motif matrices (**Figure 7B**) by choosing random frequencies for each motif and taking sets of cell types that were of total size 100 cells (for *X*_0_ and *X*_1_) or 99 cells (for *X*_2_). For the motif transformation using the motifs measured in the rat retina dataset, *e*_3_ was computed as *z*_3_ – *X*_3_*y*_3_ where *z*_3_ is the total number of A, B, and M cell types in the dataset, *X*_3_ is the empirical motif matrix based on the observed motifs, [((B,(A,B)), (A,A), (A,B), and (B,M)], and *y*_3_ is the number of occurrences of each motif within the dataset (**Figure S7A**). Similarly, *e*_4_ was computed as *z*_4_ – *X*_4_*y*_4_ where *z*_4_ is the total number of A, B, and M cell types in the dataset, *X*_4_ is the empirical motif matrix based on the observed motifs, [((B,(A,B)), (A,B), and (B,M)], and *y*_4_ is the number of occurrences of each motif within the dataset (**Figure S7B**). The total number of 113 A, B, and M cells across the entire dataset was used for ∑_i_(*z*_3_)_i_ and ∑_i_(*z*_4_)_i_ . R cells were omitted from this analysis because no rat retinal motifs contained R cells.

#### Quantification and statistical analysis

For resampling lineage trees, 10^4^ datasets were used to generate counts for each pattern across the resamples that resemble a normal distribution. For the motif transformation, 10^5^ datasets were generated to sample the possible space of cell type proportions. The p-value for all patterns in the paper was calculated by (1) determining whether the observed count is higher or lower than the average across the resamples (the null distribution), (2) counting the number of resamples that have counts at least as extreme as the observed counts, and (3) dividing this by the total number of resamples to obtain a one-sided p-value. P-values were adjusted to the total number of patterns analyzed using the Benjamini/Hochberg correction with false discovery rate (α) = 0.05 and a two-stage linear step-up procedure with estimation of the number of true hypotheses ^30, 31^.

#### Additional resources

GitHub repository: https://github.com/tranmartin45/linmo

linmo package documentation: https://tranmartin45.github.io/linmo

### Key resources table

| REAGENT or RESOURCE | SOURCE | IDENTIFIER |
| --- | --- | --- |
| Deposited data |  |  |
| Rat retina lineage trees | Gomes et al., 2011 | https://doi.org/10.1242/dev.059683 |
| Zebrafish retina lineage trees | He et al., 2012 | https://doi.org/10.1016/j.neuron.2012.06.033 |
| Mouse blastocyst lineage trees | Morris et al., 2010 | https://doi.org/10.1073/pnas.0915063107 |
| Mouse, rabbit, and monkey retina cell type proportions | Masland 2011 | https://doi.org/10.1167/iovs.10-7083 |
| Chick retina cell type proportions | Yamagata et al., 2021 | https://doi.org/10.7554/eLife.63907 |
| Software and algorithms |  |  |
| Python | Python Software Foundation | N/A |
| linmo (Python) | This paper | https://github.com/tranmartin45/linmo and https://doi.org/10.22002/htgfr-11t35 |
| Simulation and analysis code (Python) | This paper | https://github.com/tranmartin45/linmo and https://doi.org/10.22002/htgfr-11t35 |

