## Supplemental Figures for "Lineage motifs: developmental modules for control of cell type proportions"

Supplementary information

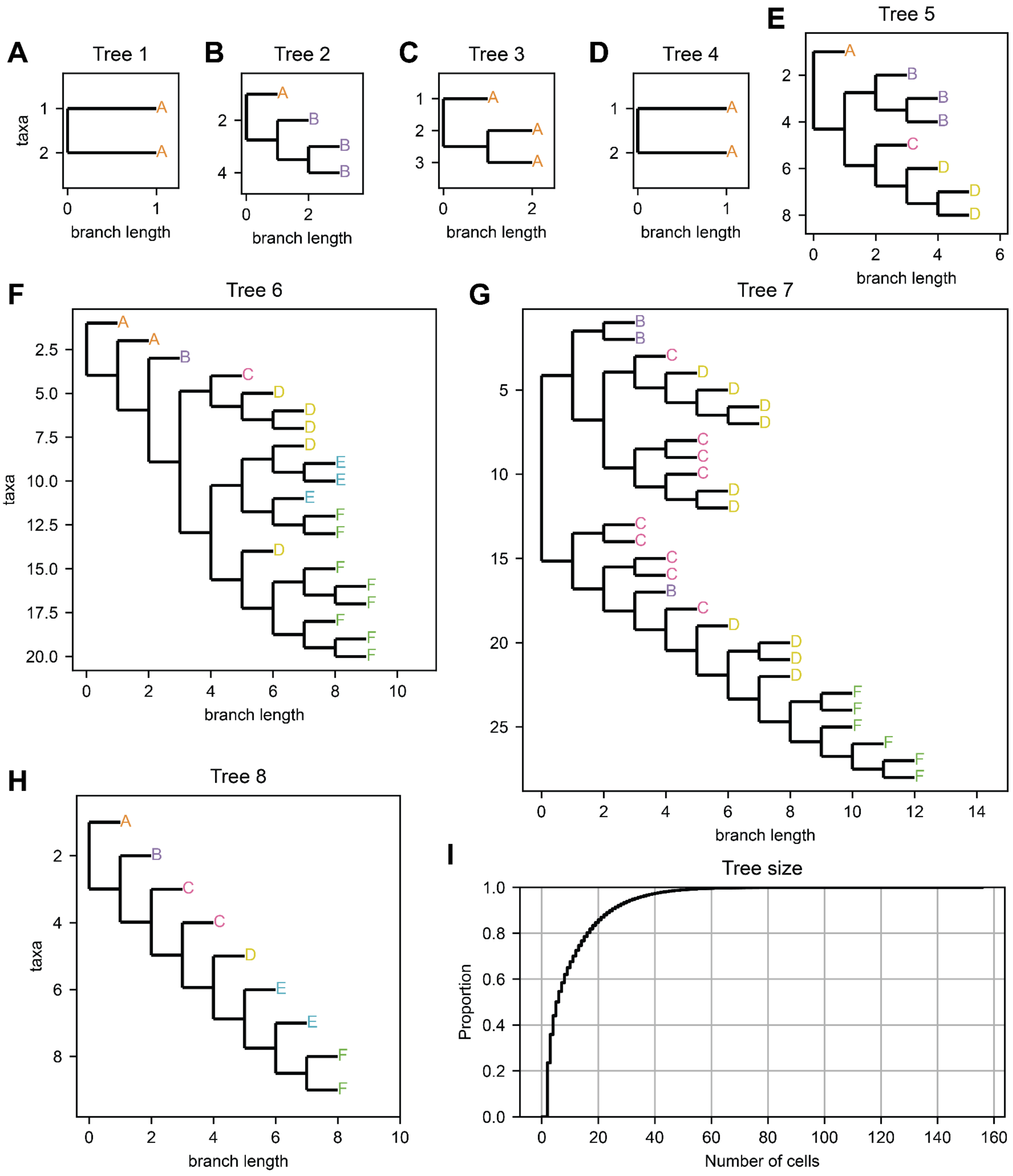


**Figure S1: Lineage trees generated based on a competence progression model of development.**

A-H. 8 random trees sampled from the simulated datasets.

I. Tree size distribution. The median sized tree contains 5 cells.

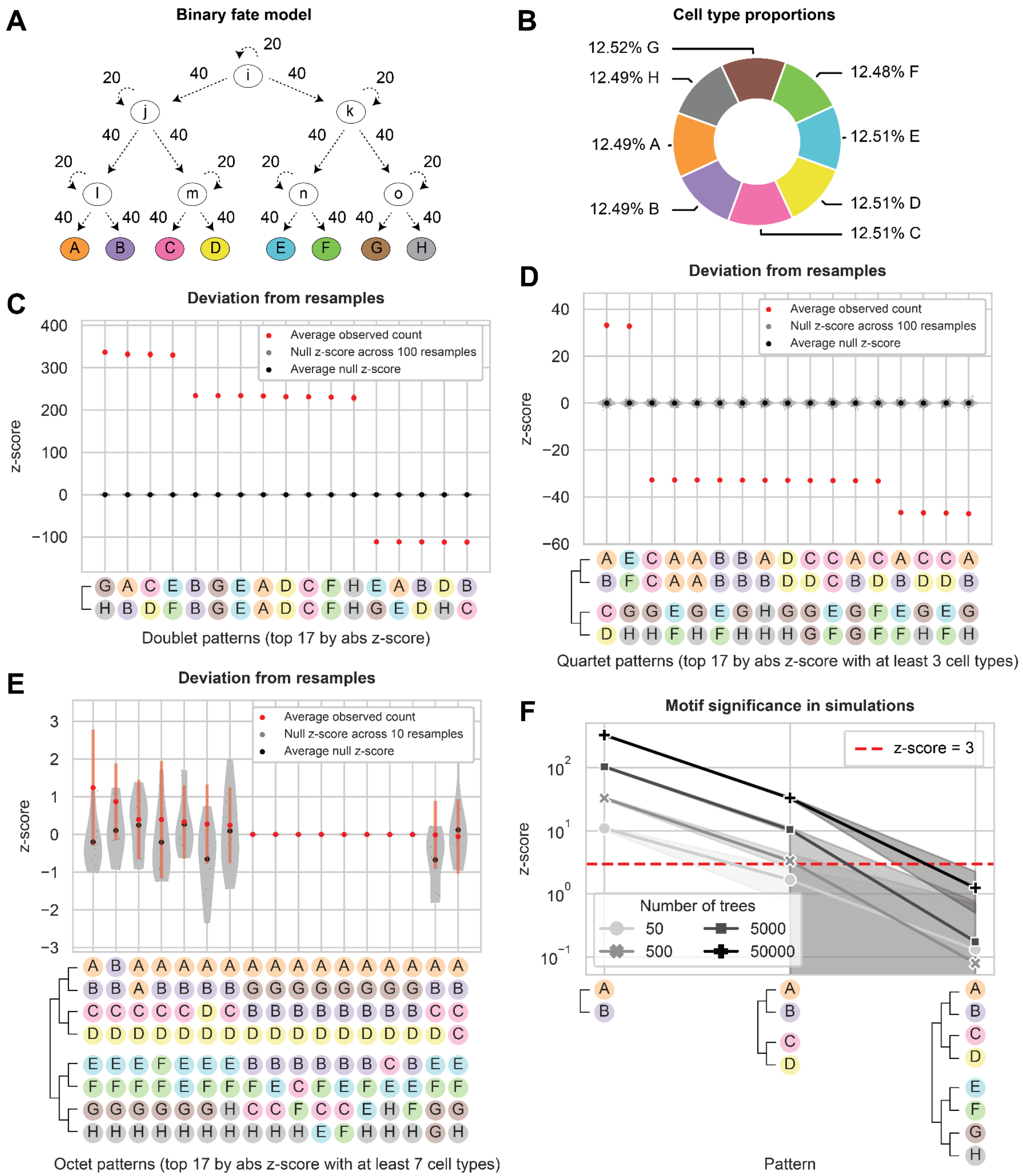


**Figure S2: Motifs reveal committed progenitors in a binary fate model of development.**

A. Lineage trees were simulated using a binary fate model of development. The upstream progenitors (‘i’, ‘j’, and ‘k’) can either self-renew with 20% probability or differentiate into two committed progenitors with 40% probability each. Downstream progenitors (‘l’, ‘m’, ‘n’, and ‘o’) can either self-renew with 20% probability or differentiate into two terminal fates with 40% probability each.

B. Cell type proportions in 500 simulated lineage trees.

C. Deviation score for top 17 most significant doublet patterns, calculated using the mean and standard deviation of counts across 10000 resamples. Null z-scores were calculated by comparing a random resample dataset to the rest of the resample datasets. 10 datasets containing 50000 simulated trees each were used, with the standard deviation across the datasets plotted as error bars.

D. Deviation score for top 17 most significant quartet patterns with at least 3 cell types.

E. Deviation score for top 17 most significant octet patterns with at least 7 cell types.

F. Deviation score for select patterns that reflect sequential differentiation of cell fates using datasets of varying size. Shading indicates 95% confidence interval across 10 datasets for each point.

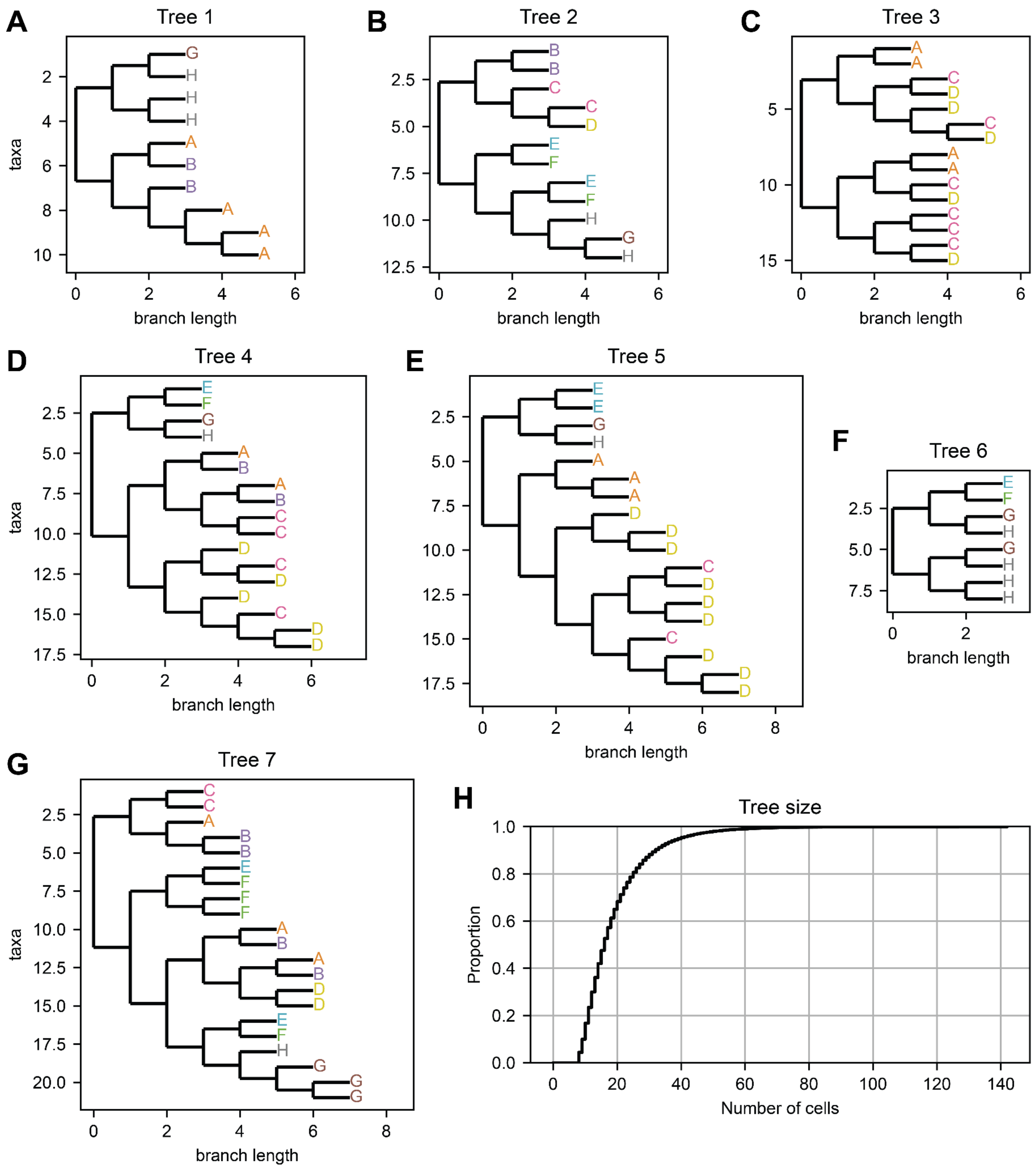


**Figure S3: Lineage trees generated based on a binary fate model of development.**

A-H. 8 random trees sampled from the simulated datasets.

I. Tree size distribution. The median sized tree contains 16 cells.

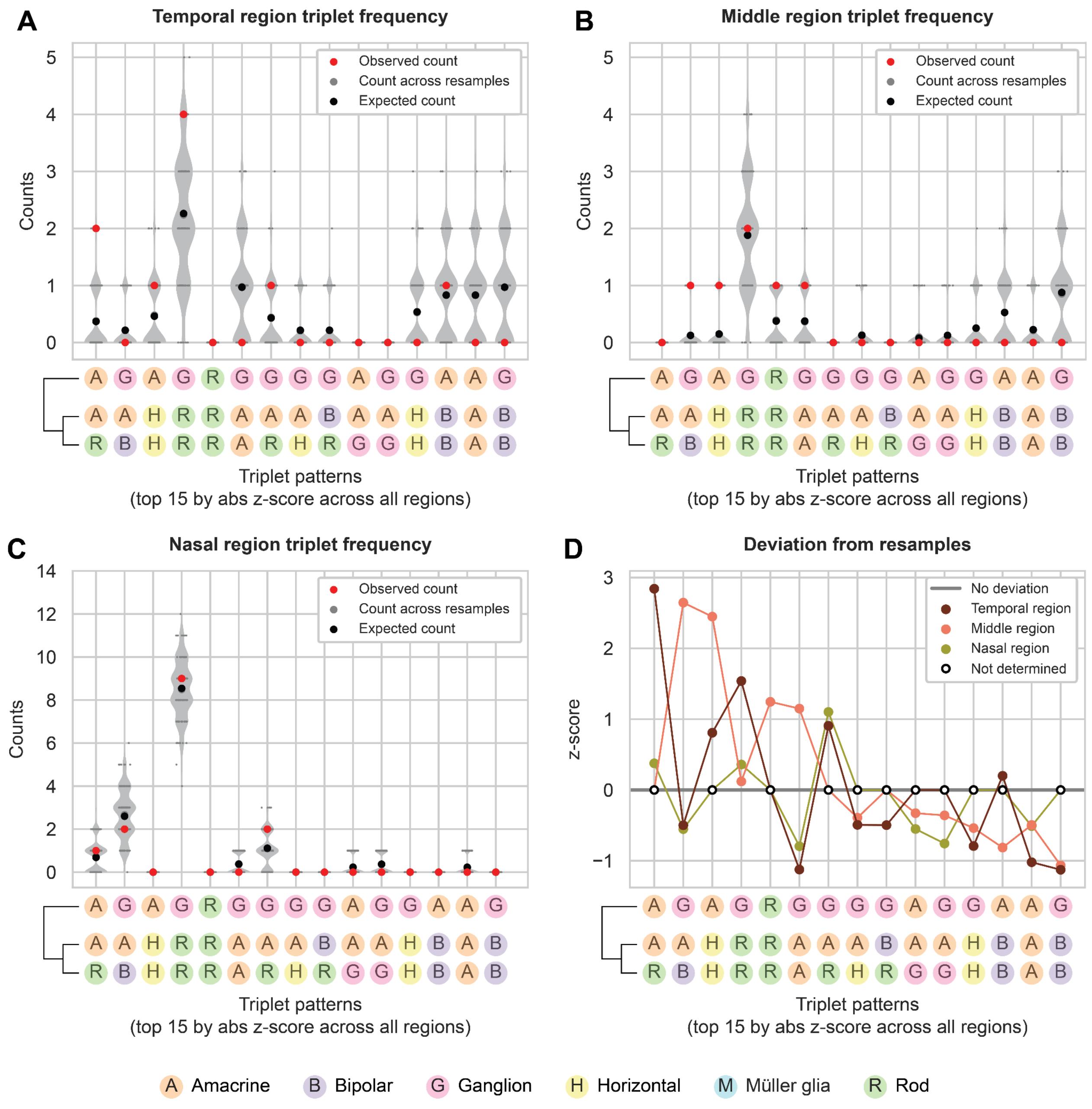


**Figure S4: No triplet patterns show significant deviation from the null expectation in the zebrafish retina dataset.**

A. Counts for triplet patterns in the observed zebrafish retina trees from He et al. [35] in the temporal region and across 10000 resamples. All 10000 resamples are represented in the violin plots, but a random subset of only 100 resamples are shown as overlaying dot plots. The top 15 significant triplets across progenitors from all spatial regions are shown. The expected count was calculated analytically (**Methods**).

B. Counts for triplet patterns in the middle region of zebrafish retina and across 10000 resamples.

C. Counts for triplet patterns in the nasal region of zebrafish retina and across 10000 resamples.

D. Deviation score for triplet patterns in the temporal, middle, and nasal region, calculated using the mean and standard deviation of counts across 10000 resamples. Triplet patterns with an observed and expected count of 0 were omitted from analysis.

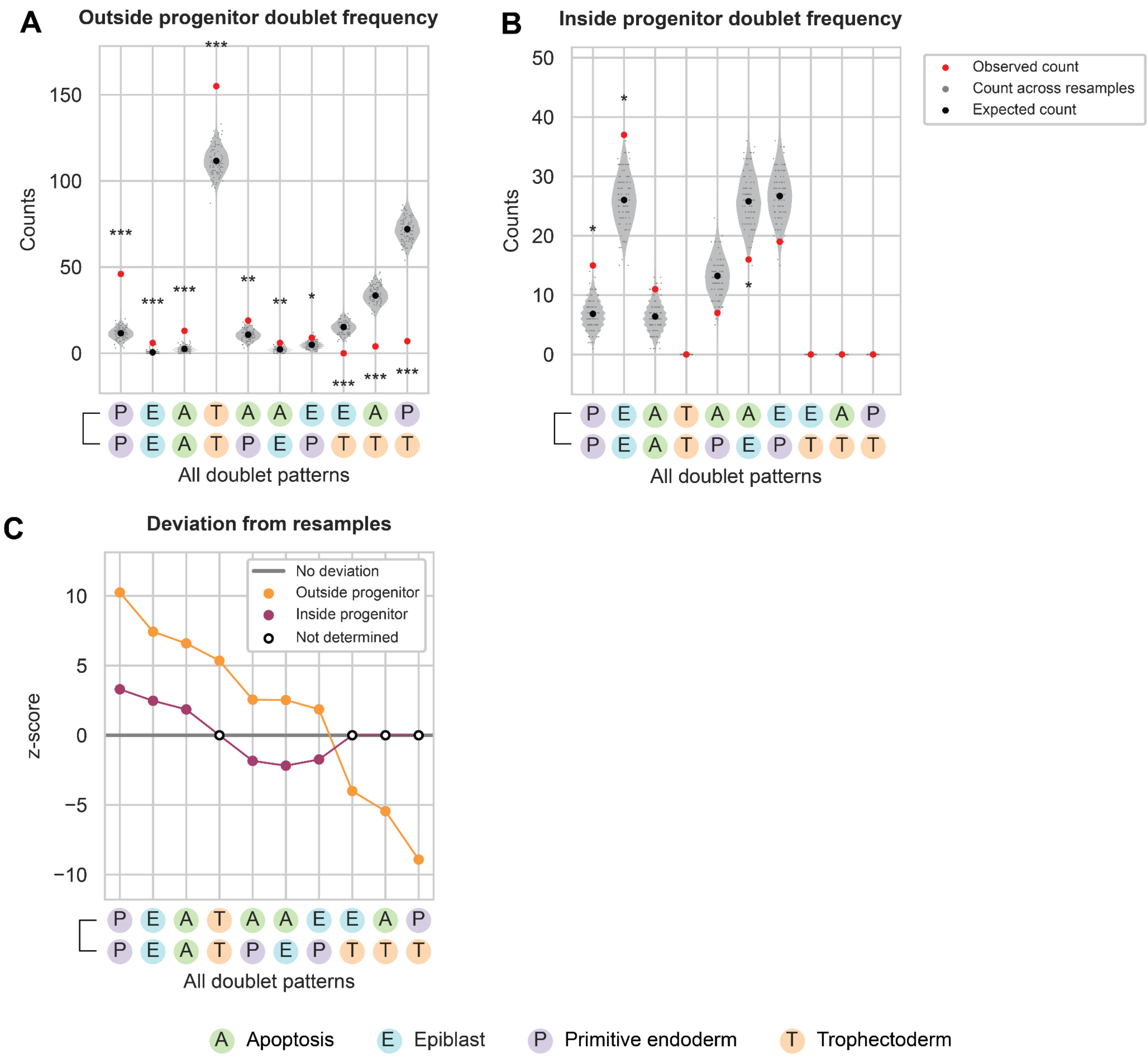


**Figure S5: Doublet lineage motif analysis in mouse blastocyst development suggests weaker fate commitment for inside progenitors at the last cell division, compared to outside progenitors.**

A. Counts for all doublet patterns in the observed mouse blastocyst trees from Morris et al. [38] in the outside progenitors and across 10000 resamples (* = adjusted p-value < 0.05; ** = adjusted p-value < 0.005, *** = adjusted p-value < 0.0005). All 10000 resamples are represented in the violin plots, but a random subset of only 100 resamples are shown as overlaying dot plots. The expected count was calculated analytically (**Methods**).

B. Counts for all doublet patterns in the observed mouse blastocyst trees in the inside progenitors and across 10000 resamples.

C. Deviation score for all doublet patterns in the outside and inside progenitors, calculated using the mean and standard deviation of counts across 10000 resamples. Doublet patterns with an observed and expected count of 0 were omitted from analysis.

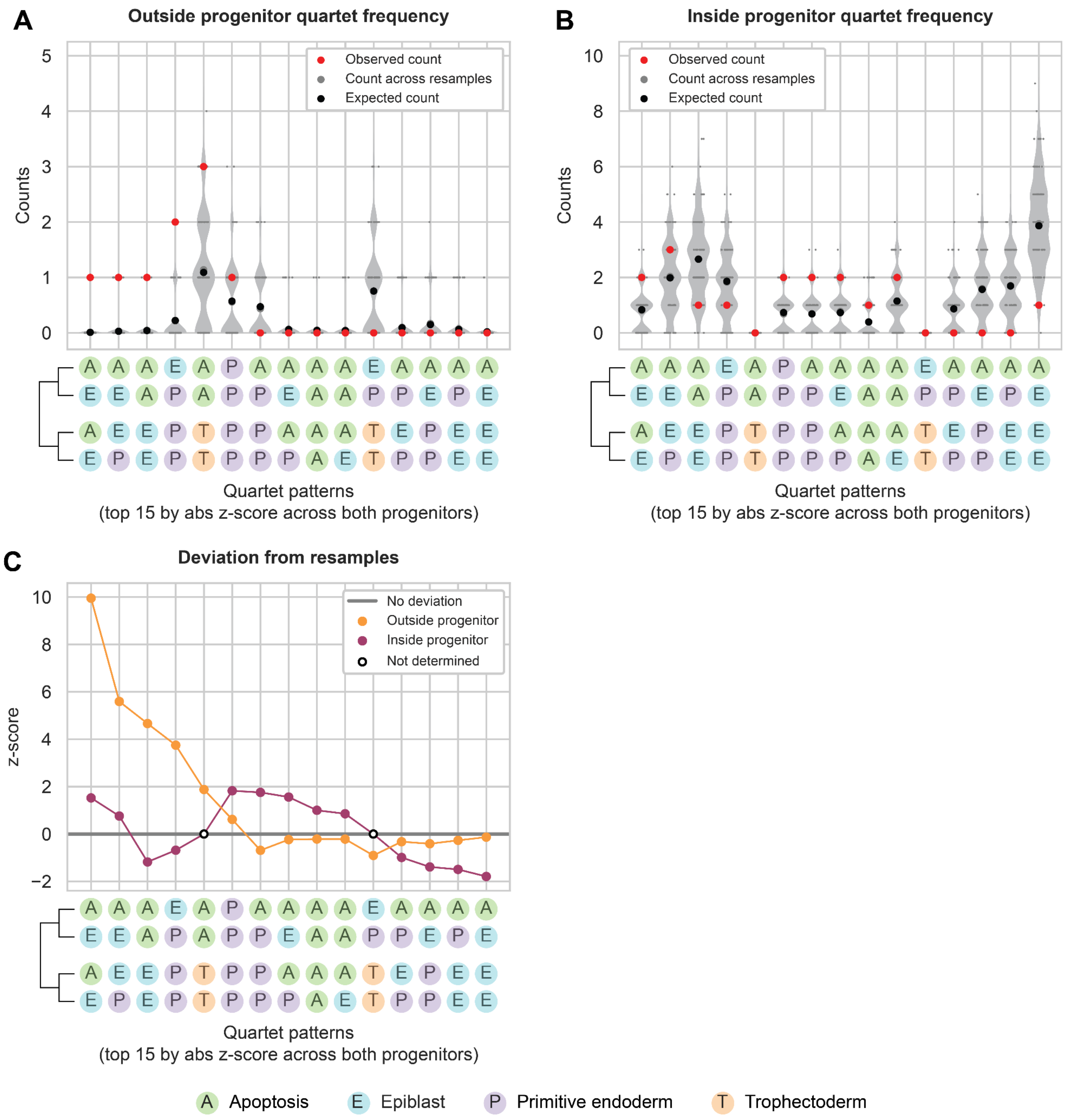


**Figure S6: No quartet patterns show significant deviation from the null expectation in the mouse blastocyst dataset.**

A. Counts for quartet patterns in the observed mouse blastocyst trees from Morris et al. [38] in the outside progenitors and across 10000 resamples. All 10000 resamples are represented in the violin plots, but a random subset of only 100 resamples are shown as overlaying dot plots. The top 15 significant quartets across both sets of progenitors are shown. The expected count was calculated analytically (**Methods**).

B. Counts for quartet patterns in the observed mouse blastocyst trees in the inside progenitors and across 10000 resamples.

C. Deviation score for quartet patterns in the outside and inside progenitors, calculated using the mean and standard deviation of counts across 10000 resamples. Quartet patterns with an observed and expected count of 0 were omitted from analysis.

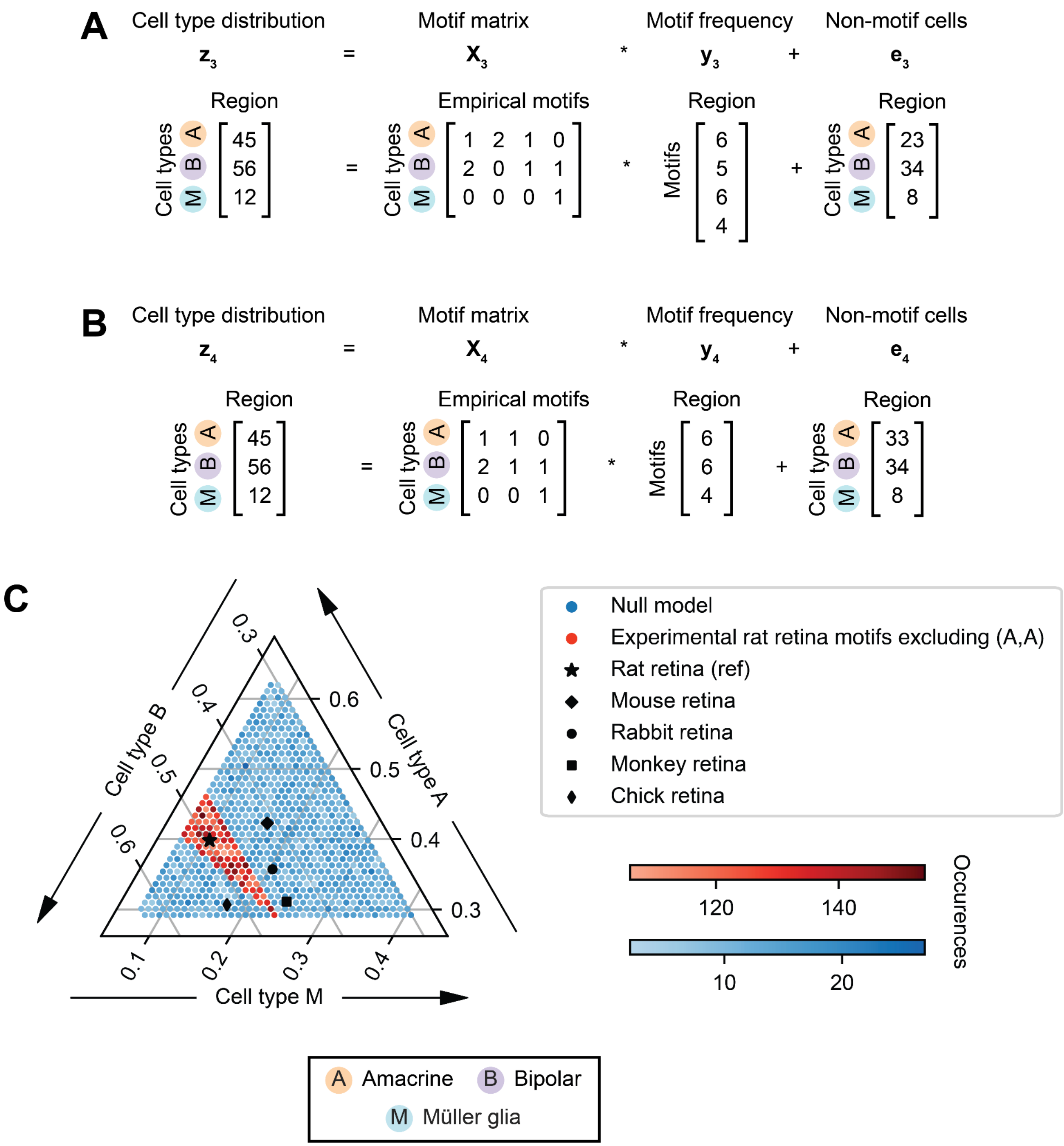


**Figure S7: The loss of the (A,A) motif reduces the subspace of accessible cell type proportions.**

A. The $z(s) = X * y(s) + e(s)$ matrix equation describes the linear transformation from motif frequencies to cell type distributions. The empirically determined values for each term, based on the rat retina dataset, are shown as $z_3 = X_3 * y_3 + e_3$.

B. The (A,A) doublet motif was omitted from the motif matrix $X_3$ and all A cells born through an (A,A) doublet were counted as part of the non-motif vector $e_3$. The perturbed model is represented as $z_4 = X_4 * y_4 + e_4$.

C. Cell type distributions were simulated using a null model ($X_0$) or the empirical motif matrix based on the rat retina motifs, excluding the (A,A) motif, which is represented as $X_4$. The lower bounds were set at $e_4$, the counts for all cell types in the experimental rat retina dataset that were born outside of a motif, including all A cells born through an (A,A) doublet. The upper bound was set by constraining the total number of cells to be the same as in the rat retina dataset, $\sum_i (z_4)_i = const$. The data was plotted as a ternary plot where each axis corresponds to the proportion of one cell type, with the cell type proportions of mouse, rabbit, monkey, and chick retina from Masland [2] and Yamagata et al. [39] overlaid.
